# A Deep Dive into the Globin Superfamily of Sharks, Skates, and Rays: Contrasting patterns of gene loss and retention relative to bony vertebrates

**DOI:** 10.1101/2025.09.19.677436

**Authors:** Hunter K. Walt, Joseph A. Hinton, Shigehiro Kuraku, Juan C. Opazo, Jay F. Storz, Federico G. Hoffmann

**Affiliations:** Department of Biochemistry, Nutrition, and Health Promotion, Mississippi State University, Mississippi State, MS 39762, USA; Molecular Life History Laboratory, Department of Genomics and Evolutionary Biology, National Institute of Genetics, Mishima, Shizuoka, Japan; Department of Genetics, Sokendai (Graduate University for Advanced Studies), Mishima, Shizuoka, Japan; Facultad de Medicina, Universidad San Sebastián, Valdivia, Chile; Integrative Biology Group, Valdivia, Chile; School of Biological Sciences, University of Nebraska, Lincoln, NE 68588, USA; Institute for Genomics, Biocomputing & Biotechnology, Mississippi State University, Mississippi State, MS 39762, USA

## Abstract

The globin gene superfamily encodes oxygen-binding proteins that are present in all domains of life. Hemoglobins and myoglobins of jawed vertebrates are among the most well-studied proteins in the context of structure-function relationships and evolution after gene duplication. However, these studies have primarily focused on bony vertebrates, and research on globin gene evolution in cartilaginous fish has been limited by a lack of genomic resources. In this study, we leverage newly available cartilaginous fish genomes to investigate globin gene family evolution across skates, rays, sharks, and sawfish. We found that, when present, most globin genes are in a single copy, with androglobin, globin-Y, and myoglobin present in all cartilaginous fish, while the two globin-X genes were differentially retained between the Holocephali and sharks, skates, and rays. Neuroglobin appears to have been lost at the common ancestor of all cartilaginous fish. The α- and β-globin gene subfamilies underwent independent expansions in different lineages of cartilaginous fish. Most cartilaginous fish globins have conserved synteny with other jawed vertebrates except myoglobin. Additionally, NPRL3, which directly flanks the hemoglobin clusters of other jawed and jawless vertebrates and regulates hemoglobin gene expression, is on a separate chromosome from the hemoglobin clusters of cartilaginous fish. When we examined globin gene expression patterns across cartilaginous fish tissues and developmental stages, we found that most globins are expressed as expected compared to other jawed vertebrates. However, hemoglobin paralogs are more widely expressed in embryonic tissues compared to later-stage tissues in cases where many copies exist. Our results reveal similar and contrasting patterns of globin gene evolution between cartilaginous and bony vertebrates and shed light on the early stages of globin gene evolution in gnathostomes.

**Significance Statement:** The evolution of the globin gene family in jawed vertebrates is of significant interest; however, most studies have been limited to bony vertebrates. With the influx of new publicly available cartilaginous fish genomes, we conducted the most comprehensive analysis of globin gene evolution in cartilaginous fish to date using phylogenetic, structural, and transcriptomic analyses. Our results shed light on similar and contrasting patterns of globin-gene evolution between cartilaginous and bony vertebrates and provide insight into the early evolution of globins in jawed vertebrates.

**Key-words:** Globins, Cartilaginous Fish, Chondrichthyes, Gene Family Evolution

## Introduction

Globins are oxygen-binding proteins present in Archaea, Bacteria, and Eukarya (Storz, 2019; Vinogradov et al., 2005). They share a similar structure of a heme prosthetic group sandwiched between α-helices, called the ‘globin fold’. The hemoglobin and myoglobin proteins of jawed vertebrates have made fundamental contributions to our understanding of structure-function relationships and the role of gene duplication in promoting functional innovation. Horse hemoglobin and whale myoglobin were the first proteins to have their crystal structures solved (Kendrew et al., 1958; Perutz et al., 1960), and sequence similarities among the human α- and β- globin chains led Ingram to propose that the subunits of tetrameric hemoglobin and myoglobin emerged via duplication and divergence of an ancestral single-copy gene (Ingram, 1961). The discovery of cyclostome hemoglobins, which are monomeric when bound to oxygen and assemble into dimers or tetramers in the deoxy state (Brittain & Wells, 1986), fit reasonably well with the scenario proposed by Ingram, with the dimeric hemoglobin of jawless vertebrates occupying an intermediate position between the proposed ancestral monomeric hemoglobin and the tetrameric hemoglobins of jawed vertebrates (Goodman et al., 1975; Müller et al., 2003).

Up until the end of the 20^th^ century, myoglobin, α-hemoglobin and β-hemoglobin of jawed vertebrates and the hemoglobins and myoglobins of jawless fish were the only known vertebrate globins. The advent of comparative genomics resulted in a new appreciation of the diversity of the globin gene superfamily in the animal kingdom (Burmester et al., 2000, 2002; Fuchs et al., 2006; Hoffmann et al., 2010; Hoffmann, Opazo, Hoogewijs, et al., 2012; Kugelstadt et al., 2004; Opazo, Lee, et al., 2015; Roesner et al., 2005). Phylogenetic and synteny analyses revealed that vertebrate globins diversified via a combination of tandem gene duplications and whole-genome duplications (Hoffmann et al., 2021; Hoffmann, Opazo, & Storz, 2012; Opazo et al., 2013; Storz et al., 2013). Currently, the globins of vertebrates can be separated into four groups: Androglobin (Adgb), Neuroglobin (Ngb), globin-Xs (GbX1 and GbX2), and the vertebrate-specific globins. The latter group includes cytoglobin (Cygb), globin-E (GbE), globin-Y (GbY), the myoglobin (Mb), α-globins (HbA) and β-globins (HbB) of gnathostomes (jawed vertebrates) and the hemoglobins (cHb) and myoglobins (cMb) of cyclostomes (jawless vertebrates).

Many of these newly discovered globins do not have canonical oxygen-transport or oxygen-storage functions, functions that evolved independently in gnathostomes and cyclostomes, respectively (Hoffmann et al., 2010; Schwarze et al., 2014). The pro-ortholog of the cMb and cHb genes of cyclostomes is closely related to Cygb, whereas the HbA and HbB gene families of gnathostomes are more closely related to Mb and GbE. The last common ancestor of vertebrates included copies of Adgb, GbX1, GbX2, Cygb, Ngb, GbY, the preduplication progenitor of the GbE and Mb genes, the progenitor of the α- and β-globin gene families of gnathostomes, and the progenitor of the cMb and cHb genes of cyclostomes. Lineage-specific duplications and deletions of members of this ancestral gene set account for the observed variation in the globin gene repertoires among contemporary vertebrates.

The early stages of globin evolution in gnathostomes have received significant interest and remain poorly understood. Morris Goodman and Motoo Kimura argued about the role of Darwinian selection on the early divergence between Mb and Hb (Goodman, 1981; Goodman et al., 1975; Kimura, 1981), and the branching of vertebrate-specific globins has proven hard to resolve. This is particularly true for the globin gene repertoire of cartilaginous fishes (Class Chondrichthyes), largely due to a relative dearth of genomic sequence data in comparison to resources available for bony vertebrates. The split between bony fishes and cartilaginous fishes is the deepest in the tree of extant jawed vertebrates, dating back to ∼ 460 million years ago (mya). Within extant cartilaginous fishes, the split between Elasmobranchii (sharks and batoids) and Holocephali (chimaeras and ratfishes) is the deepest, timed to ∼415 mya, whereas the split between sharks, superorder Selachimorpha; and batoids, which comprise rays, skates, torpedoes, and sawfish in the superorder Batoidea, is timed to ∼270 mya. Until recently, research into the globin repertoire of cartilaginous fish had been limited to the elephant fish, *Callorhinchus milii*, a representative of the subclass Holocephali that has one of the smallest globin repertoires among vertebrates (Opazo, Lee, et al., 2015). After a long period with no additional genomic resources, chromosome-level assemblies of elasmobranch genomes are now available for diverse species, which presents an excellent opportunity to expand our understanding of the evolution of the globin gene family in this group.

Here we combine bioinformatic searches with phylogenetic and synteny analyses to characterize the globin repertoires in the genomes of sharks, rays, skates and sawfish. We also take advantage of available transcriptome data to compare patterns of gene expression across multiple tissues and developmental stages. We report new findings regarding the differential retention of ancestral globin genes in cartilaginous fishes relative to bony vertebrates.

## Results and Discussion

### Bioinformatic Searches

We identified a total of 261 globin sequences from 22 species of cartilaginous fish, 22 of which were Adgbs (fig. 1, Table 1, Supplementary Table 1, Supplementary Material online). Our searches revealed that almost all genomes screened had single copies of Adgb, GbY, and Mb, that most genomes possessed multiple copies of α- and β-globin, and that Cygb was present only in horn shark and elephant fish (fig. 1). Our initial searches were limited to 20 RefSeq genomes (Table 1), where we identified 243 previously annotated globins. To this data set, we added the Adgb and Cygb genes of the white shark, the GbY gene of the red stingray, the GbX1 of the smalltooth sawfish, and the globin repertoires of the rabbitfish and the ratfish using manual annotations. Even though the Adgbs of white shark, ratfish, and rabbitfish were found in the corresponding genomes, the hits cover >75% of the translated protein and are clearly incomplete. We infer that these genes are present and functional, but not fully covered in the current genome releases.

**Figure 1:**
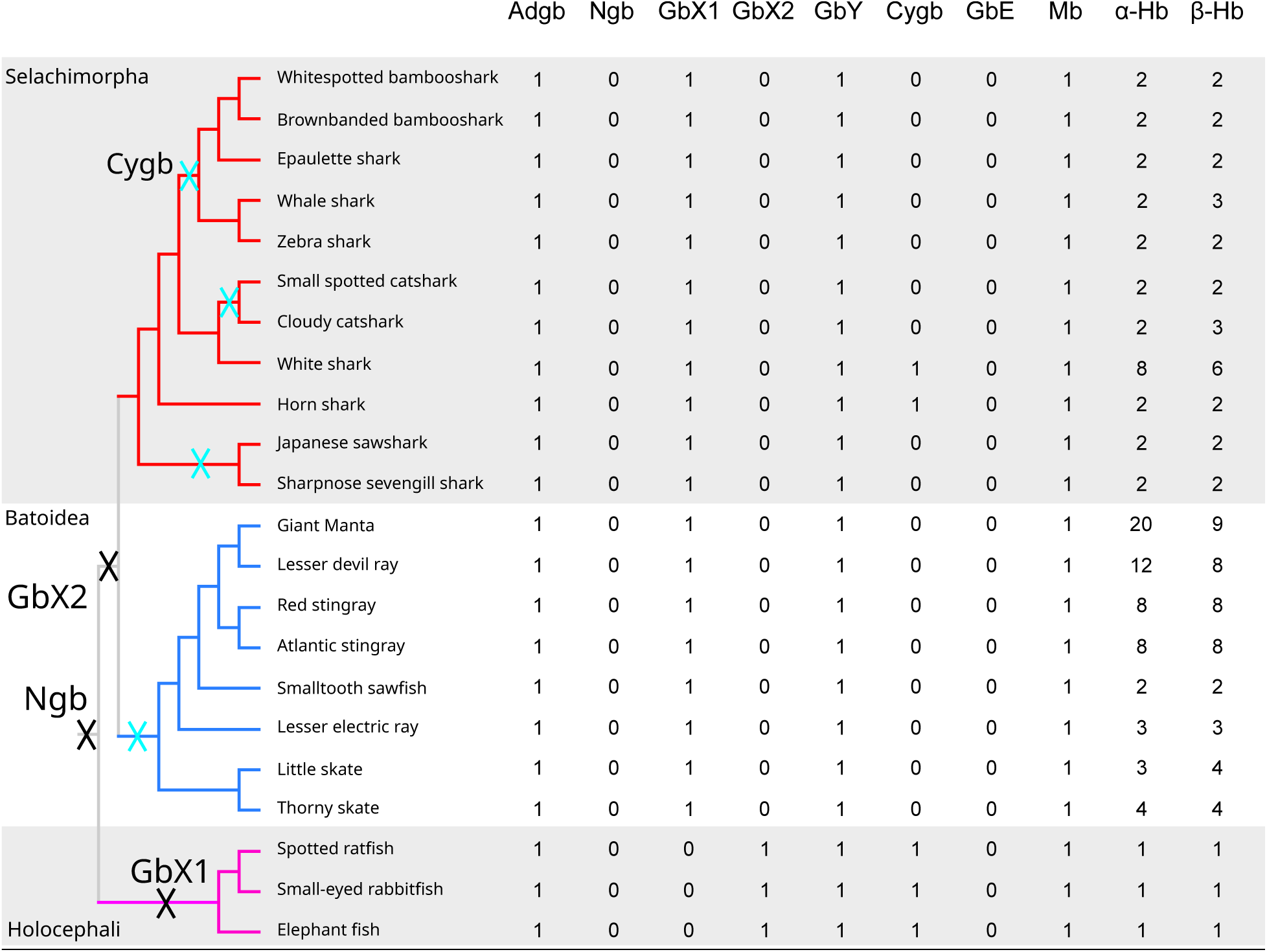
Phyletic distribution of the different globin subfamilies in the genomes of the cartilaginous fishes in our study. The counts correspond to the number of globin genes after data curation. Xs along the tree correspond to inferred losses Ngb, GbX1, GbX2, and Cygb (cyan).

**Table 1.** Cartilaginous fish genomes surveyed in the study.

| RefSeq | Common Name | Species Name | Superorder | Order | Accession |
| --- | --- | --- | --- | --- | --- |
| 1 | Whitespotted bambooshark | *Chiloscyllium plagiosum* | Selachimorpha | Orectolobiformes | GCF_004010195.1 |
| 1 | Brownbanded bambooshark | *Chiloscyllium punctatum* | Selachimorpha | Orectolobiformes | GCF_047496795.1 |
| 1 | Epaulette shark | *Hemiscyllium ocellatum* | Selachimorpha | Orectolobiformes | GCF_020745735.1 |
| 1 | Whale shark | *Rhincodon typus* | Selachimorpha | Orectolobiformes | GCF_021869965.1 |
| 1 | Zebra shark | *Stegostoma tigrinum* | Selachimorpha | Orectolobiformes | GCF_030684315.1 |
| 1 | Horn shark | *Heterodontus francisci* | Selachimorpha | Heterodontidae | GCF_036365525.1 |
| 1 | Smaller spotted catshark | *Scyliorhinus canicula* | Selachimorpha | Carcharhiniformes | GCF_902713615.1 |
| 1 | Cloudy catshark | *Scyliorhinus torazame* | Selachimorpha | Carcharhiniformes | GCF_047496885.1 |
| 1 | Great white shark | *Carcharodon carcharias* | Selachimorpha | Carcharhiniformes | GCF_017639515.1 |
| 1 | Japanese sawshark | *Pristiophorus japonicus* | Selachimorpha | Pristiophoriformes | GCF_044704955.1 |
| 1 | Sharpnose sevengill shark | *Heptanchias perlo* | Selachimorpha | Hexanchiformes | GCF_035084215.1 |
| 1 | Giant Manta | *Mobula birostris* | Batomorphi | Myliobatiformes | GCF_030028105.1 |
| 1 | Lesser devil ray | *Mobula hypostoma* | Batomorphi | Myliobatiformes | GCF_963921235.1 |
| 1 | Red stingray | *Hemitrygon akajei* | Batomorphi | Myliobatiformes | GCA_048418815.1 |
| 1 | Atlantic stingray | *Hypanus sabinus* | Batomorphi | Myliobatiformes | GCF_030144855.1 |
| 1 | Smalltooth sawfish | *Pristis pectinata* | Batomorphi | Rhinopristiformes | GCF_009764475.1 |
| 1 | Lesser electric ray | *Narcine bancroftii* | Batomorphi | Torpediniformes | GCF_036971445.1 |
| 1 | Little skate | *Leucoraja erinaceus* | Batomorphi | Rajiformes | GCF_028641065.1 |
| 1 | Thorny skate | *Amblyraja radiata* | Batomorphi | Rajiformes | GCF_010909765.2 |
| 0 | Spotted ratfish | *Hydrolagus colliei* | Holocephali | Holocephali | GCA_035084275.1 |
| 0 | Small-eyed rabbitfish | *Hydrolagus affinis* | Holocephali | Holocephali | GCA_012026655.1 |
| 1 | Elephant fish | *Callorhinchus milii* | Holocephali | Holocephali | GCF_018977255.1 |

### Data Curation

There are two GbX genes in the cloudy catshark genome, NCBI Gene IDs 140406215 and 140406187, both in short contigs that do not include any additional annotated genes. These two genes are predicted to encode proteins of 152 and 136 amino acids that are identical over the positions they share. We infer that they are alleles rather than duplicate genes, and we retained the more complete sequence for downstream analyses. In the horn shark we found two Cygb genes, NCBI Gene IDs 137366896 and 137359322, which encode identical proteins. The 137366896 gene is in a contig without flanking genes, whereas the 137359322 gene is in the expected genomic location. As with the GbX duplicates of the Cloudy catshark, we assumed that these genes are alleles rather than duplicates, so we retained the sequence of the 137359322 gene for all subsequent analyses. As in Opazo et al. (2015), we did not find matches for two elephant fish transcripts identified in a Sanger-sequencing study of full-length cDNAs (Tan et al., 2012). For the sake of consistency, these genes were not included in the counts, but they are included in the phylogeny to enable comparisons with the previous study.

### Globin Repertoires

In our curated set, the number of globins per genome ranges from 7 in the three representatives of Holocephali (elephant fish, rabbitfish and ratfish), to 34 in the giant manta (fig. 1). Variation in the number of globins per genome is largely driven by changes in the α- and β-globin gene subfamilies, which range from 1 to 20 and from 1 to 9 copies per genome respectively. Cygb was absent in most cartilaginous fish, while the Adgb, GbY, GbX, and Mb genes were present as single copies in all taxa. As in our previous report (Opazo, Lee, et al., 2015), we did not find traces of either Ngb or GbE in any of the cartilaginous fish genomes we screened.

### Phylogenetic Analyses

We conducted separate phylogenetic analyses for Adgb and all other globins combined. In addition, we conducted separate analyses for each globin subfamily where the arrangement of cartilaginous fish sequences deviated from the expected organismal relationships among Holocephali, Batoidea, and Selachimorpha. To provide evolutionary context, we included additional representatives of deuterostomes as well as bony vertebrates in all phylogenetic analyses. The estimated phylogeny of Adgb sequences (fig. 2) matches the organismal tree (fig. 1), with the Adgb genes of Selachimorpha, Batoidea, and Holocephali forming reciprocally monophyletic groups, and Selachimorpha and Batoidea placed sister to one another.

**Figure 2:**
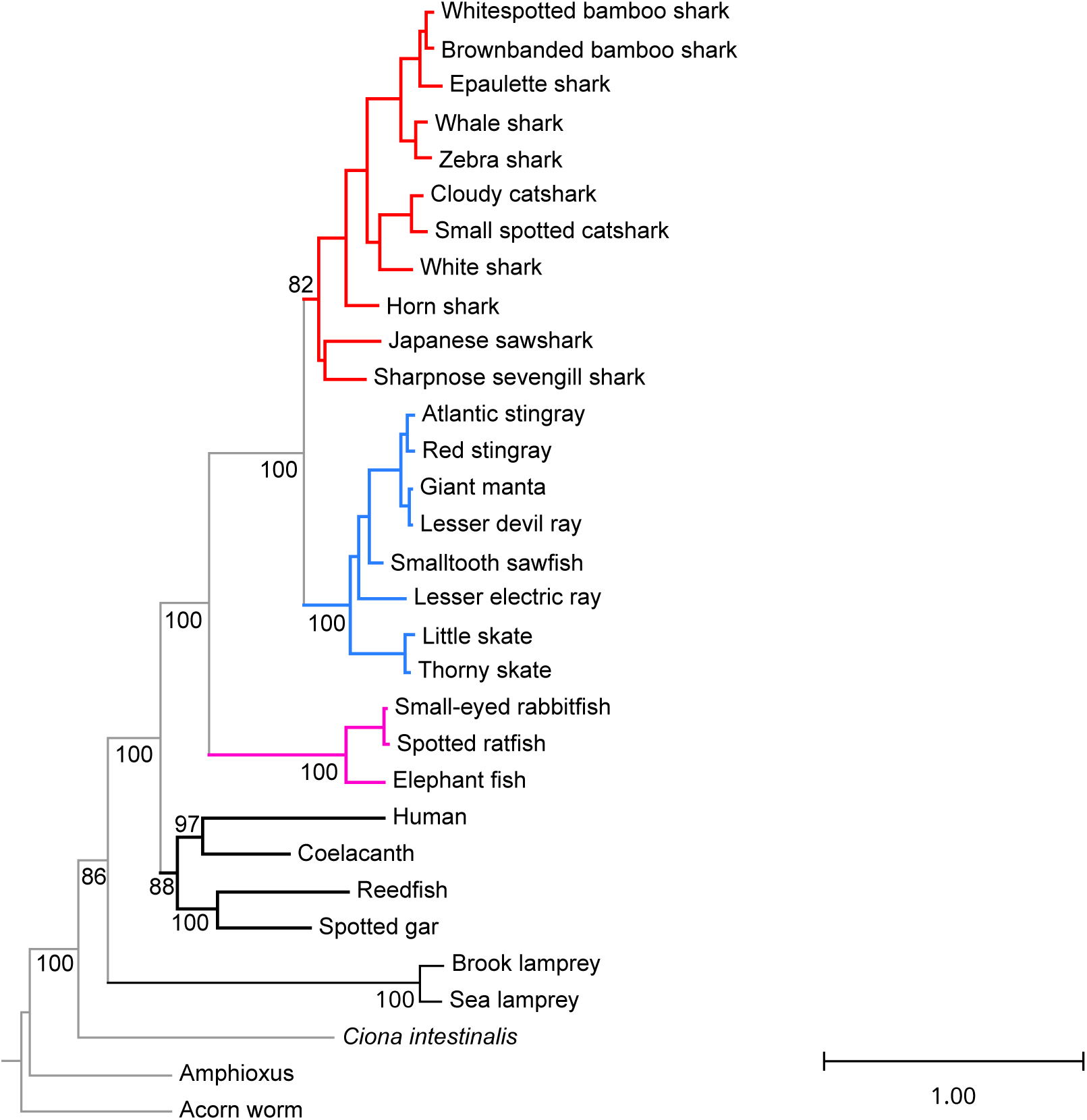
Maximum likelihood phylogram depicting relationships among the Androglobin genes of cartilaginous fishes, with bony vertebrate and deuterostome sequences included for context. Numbers correspond to ultrafast bootstrap values. Shark branches are in red, batoid branches are in blue, and holostei branches are in fuchsia.

In the case of all other globins other than Adgb, and in agreement with previous studies (Hoffmann, Opazo, & Storz, 2012; Opazo, Lee, et al., 2015; Prothmann et al., 2020), the globins of vertebrates fell into three separate clades (fig. 3, Supplementary fig. 1, Supplementary Material online), with Neuroglobin in one, Globin X in the second, and vertebrate specific globins in the third, with representatives of cartilaginous fishes in the latter two groups. The monophyly of the clade of vertebrate GbXs clade and of the clade of vertebrate specific globins were relatively high, 99% and 88% respectively, and the monophyly of each vertebrate specific globin subfamily had strong support as well, with bootstrap values of 99 or 100%, with the exception of GbX1, which had a bootstrap support value of 68% (Figs. 2 and 3, Supplementary fig. 1, Supplementary Material online).

**Figure 3:**
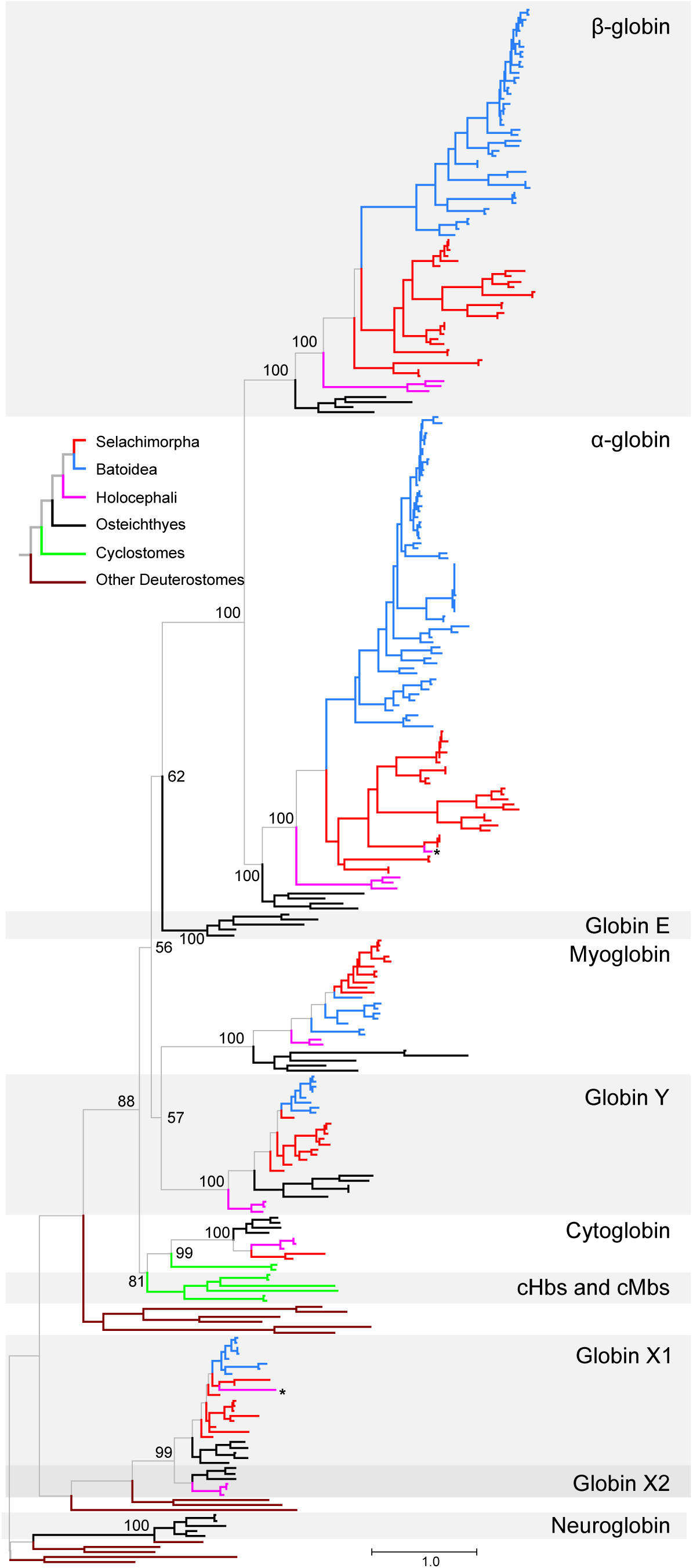
Maximum likelihood phylogram depicting relationships among the single-domain globin genes of cartilaginous fishes, with bony vertebrate and deuterostome sequences included for context. Numbers correspond to ultrafast bootstrap values. Shark branches are in red, batoid branches are in blue, and holostei branches are in fuchsia. Genes marked with asterisks are included in the phylogeny for comparative purposes, but they are not found in the current genome assemblies. A version of this tree with terminal branches labeled is presented as Supplementary fig. 1, Supplementary Material online).

Within GbXs, we found GbX1 copies in Selachimorpha and Batoidea, and GbX2 copies in the Holocephali (fig. 3, Supplementary fig. 1 and Supplementary Table 1, Supplementary Material online). As in Opazo et al. 2015, we failed to find the elephant fish GbX1 gene in the current assembly and did not find any matches in the other two Holocephali either. Both GbX1 and GbX2 sequences are placed in monophyletic groups, with the sequences of bony vertebrates placed sister to those of cartilaginous fishes in both cases. In the case of GbX1, sequences from Batoidea were placed in a monophyletic group, whereas the GbX1 sequences from Selachimorpha were paraphyletic relative to the Batoidea. Interestingly, the elephant fish GbX1 is placed sister to the GbX1 sequences of catsharks. Forcing the cartilaginous fish GbX1s of Batoidea and Selachimorpha to be monophyletic and the elephant fish GbX1 as sister to all other cartilaginous fish GbX1s did not result in a significant loss in likelihood score (Supplementary Table 2, Supplementary fig. 2, Supplementary Material online). We only found GbX2 copies in the Holocephali, and these sequences are placed in a monophyletic group sister to the GbX2s from bony vertebrates. Thus, the single-copy GbX genes of cartilaginous fish represent a case of hidden paralogy (Kuraku, 2010), resulting from reciprocal losses of alternative GbX paralogs, GbX1 in the case of Holocephali, and GbX2 in the case of Elasmobranchs.

Within vertebrate-specific globins, the α- and β-globin genes of jawed vertebrates are in sister monophyletic groups (fig. 3, Supplementary fig. 1, Supplementary Material online).

Osteichthyes GbE is placed sister to the α- and β-globin clade, and this clade is sister to the group that unites Mb and GbY with the clade that includes Cygb and the cHbs and cMbs of cyclostomes at the deepest split of vertebrate specific globins (fig. 3, Supplementary fig. 1, Supplementary Material online). Support for the nodes resolving relationships among the different subfamilies of vertebrate specific globins is relatively low, ranging from 57 to 62%, with the exception of the sister relationship between the α- and β-globins of jawed vertebrates (bootstrap support of 100%), and the sister relationship between Cygb and the clade that unites cHbs and cMbs (bootstrap support of 81%). Within these paralogs, cartilaginous fish sequences fell in monophyletic groups in the cases of α- and β-globin, Cygb, and Mb, and the sequences for all three of Batoidea, Selachimorpha, and Holocephali fell in monophyletic groups in the α- globin subtree. The Holocephali paralogs always fell in monophyletic groups, while the Batoidea and Selachimorpha sequences did so in most cases. As in the case of GbX, we ran separate searches for the β-globin, GbY, and Mb paralogs to test whether forcing cartilaginous fish sequences to be monophyletic and, within them, forcing each of Holocephali, Batoidea, and Selachimorpha sequences to be monophyletic as well. Topology tests indicate that GbY was the only case where the constrained tree was significantly different from the maximum likelihood gene tree (Supplementary Table 2, Supplementary figs. 3-5, Supplementary Material online). A strict reconciliation of the GbY tree with the organismal tree would imply that many of these separate GbY clades derive from different ancestral genes. Since GbY has only been found as a single-copy gene when present, we infer that these genes are orthologs and attribute this unexpected phylogenetic arrangement to an unusual accumulation of substitutions in the stem of the GbY tree, as it minimizes duplication events.

As for most other vertebrate groups, the only globin paralogs to exhibit large differences in gene copy number are α- and β-globins. The three Holocephali species have a single copy of each gene, grouped in strongly supported monophyletic clades that are placed sister to the elasmobranch sequences. Within elasmobranchs, sharks and batoids exhibit contrasting patterns of evolution in the α- and β-globins. The α- and β-globin genes of sharks are arranged in species-specific clades, which suggests either a rapid rate of gene turnover or gene conversion. By contrast, batoid duplicates exhibit a more complex pattern, with both species-specific expansions, such as a 10-gene expansion of α-globin genes in the giant manta, and multiple α- and β-globin duplications shared between the Giant manta and the Lesser devil ray (Supplementary fig. 1, Supplementary Material online).

### Synteny Analyses

In most cartilaginous fish, Adgb is flanked by STXBP5 and SASH1A on one side, and by RAB32 and GRM1 on the other side. This arrangement is shared with human, spotted gar, and lamprey. Mb synteny is less well-conserved. In the elephant fish, Mb is flanked by FOXRED2 and LUC7l2 on one side and SMDT1B and NDUFA6 on the other. In sharks and batoids, it is flanked by NAGA and SERHL on one side and by LUC7L2 and DDX59 on the other. This arrangement is not shared with bony fishes. In elephant fish, Cygb is flanked by QRICH2 and RHBDF2 upstream and by PRPSAP1 and RNF157 downstream. Cygb is flanked by PRPSAP1, QRICH2, and RNF157 downstream in the horn shark and white shark, and by UBE2O and AANAT upstream in the white shark. This information is lacking for the horn shark Cygb as the gene is at the beginning of a scaffold. In human and spotted gar, the orthologs of PRPSAP1, RNF157, QRICH2, RHBDF2, UBE2O, and AANAT are all on the same chromosome as the Cygb gene. The GbY gene is located at one end of the cluster of the α- and β-globin genes. The α-, β-, and GbY globin cluster is flanked by AANAT2 and RHBDF1 on one end, and by LUC7L on the other end in the vast majority of cartilaginous fishes. Unlike most other vertebrate lineages, NPRL3, a gene which is normally located immediately adjacent to the multigene hemoglobin clusters of lampreys and bony vertebrates and contains cis-acting regulatory elements in its introns (references), is found on a separate chromosome (Supplementary fig. 6, Supplementary Material online). In mammals, the regulatory elements found within NPRL3 drive developmental changes in the expression of the α-globin genes in the adjacent cluster in the precursors of red blood cells (Miyata et al., 2020; Preston et al., 2025).

### Structural Considerations of α- and β-globins

The β-chain Hbs of sharks and batoids differ from the globins of most vertebrates in lacking a portion of the D-helix (Chong et al., 1999; Fisher et al., 1977; Naoi et al., 2001; Verde et al., 2005). A visual inspection of the β-globin alignment revealed that all cartilaginous fish sequences share the deletion of the D helix, positions 54-57 in the β-globin alignment or 164-167 in the full alignment, and that Orectolobiform and Carcharhiniform sharks have a second deletion in the same region that involves the first three amino acids of the E Helix, positions 58-60 in the β-globin alignment or 168-171 in the full alignment. The first of these deletions maps to the stem of the tree of cartilaginous fishes, and the second one to the last common ancestor of Orectolobiform and Carcharhiniform sharks. Vertebrate α-globins have a deletion in the same location, and structural predictions indicate that the β-globins of cartilaginous fish are structurally more similar to human α-globin than to human β-globin (fig. 4). The D helix appears to be present in all other sequences in our full alignment, except for two GbE proteins from lungfish and reedfish. Thus, our results indicate that the α-globins of all jawed vertebrates, the β- globins of cartilaginous fish, and the two GbE listed above convergently lost this helix. The functional relevance of these deletions is unclear, as deleting the D helix in the β-globin chains of human hemoglobins has minimal impact on the oxygen binding affinity of the tetramer (Komiyama et al., 1991).

**Figure 4:**
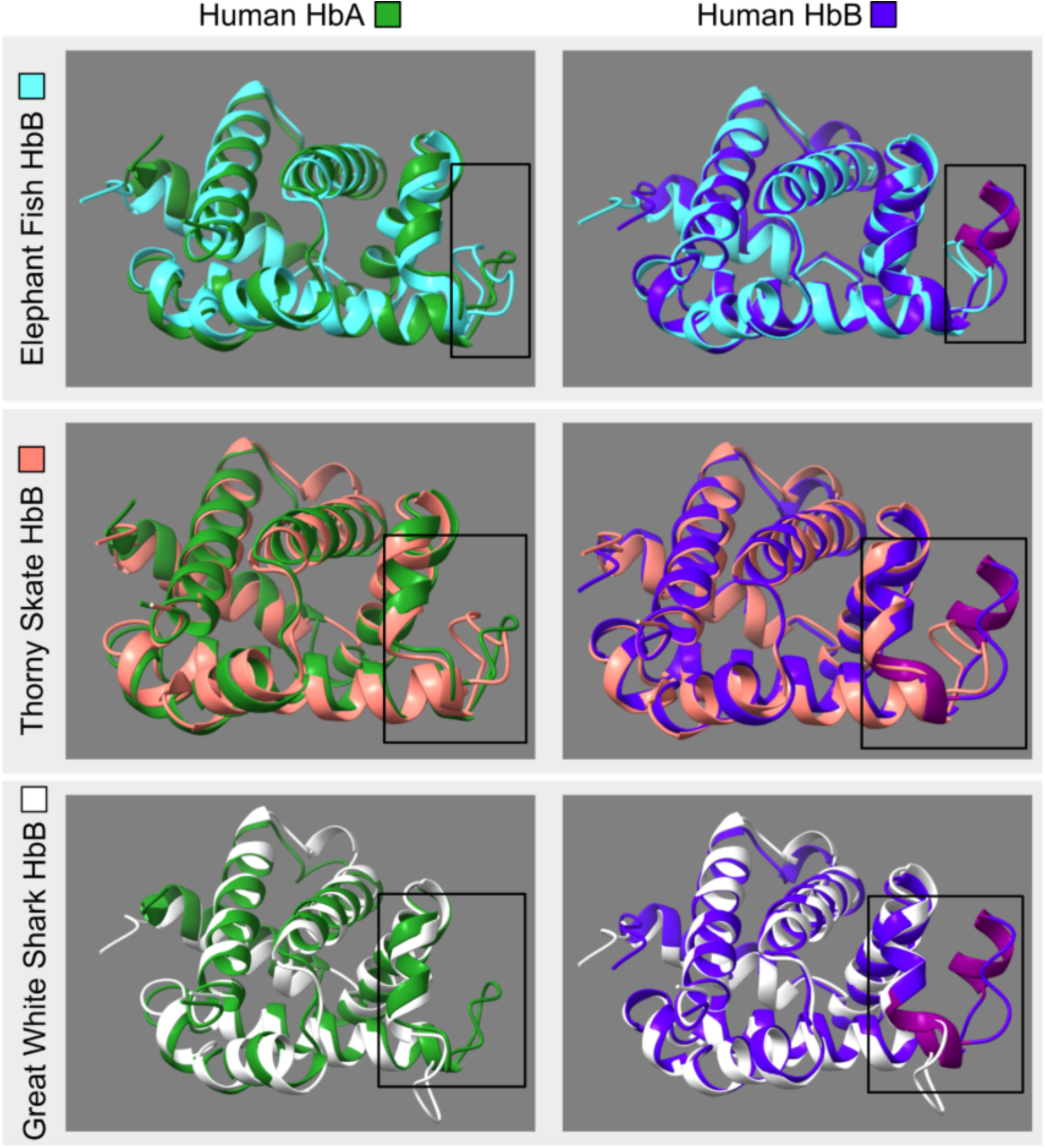
Structural Comparison of human and cartilaginous fish Hb AlphaFold Models. The predicted structures of α- and β-globin subunits from cartilaginous fish on top of human α- and β- subunits. The boxed area highlights the helix on the human β- globin subunits which is not found on the β-globin subunits from cartilaginous fishes. Accession numbers are as follows: P69905 and P68871 for human α- and β-globin, for elephant fish β-globin: NP_00127970, thorny skate β-globin: XP_032896874, and great white shark β-globin: XP_041061644.

### Gene Expression

We screened 689 RNA-Seq libraries from cartilaginous fish to assess the expression profiles of the different globin paralogs. As expected, we found that the Hb and Mb genes were the most highly expressed in all species (fig. 5, Supplementary fig. 7, Supplementary Material online). Myoglobin is most heavily expressed in the heart, whereas the α- and β-globin genes are most heavily expressed in blood. While Adgb had high expression in the brain, kidney, and various embryonic tissues, it was also highly expressed in the testis (when testis data were present), consistent with observations in mammalian tissues (Hoogewijs et al., 2012; Keppner et al., 2022). GbX1 was broadly expressed, with the highest expression in neural tissues and the eye, consistent with previous findings (Fuchs et al., 2006; Gallagher & Macqueen, 2017; Hoffmann et al., 2021; Opazo, Lee, et al., 2015). GbY is expressed across many tissues; however, we observed the highest expression in the skin, brain, gill, spleen, kidney, and intestine for the various taxa used in our study. This is consistent with previous studies (Gallagher & Macqueen, 2017; Opazo, Lee, et al., 2015; Schwarze et al., 2015), and the function of GbY remains unclear (Keppner et al., 2020). Cygb was only present in a few species – the horn shark, the elephant fish, and the white shark. Of these, the elephant fish has low levels of Cygb expression in the kidney, neural tissues, and reproductive tissues, while the Cygbs of the horn shark and white shark are not expressed (fig. 5, Supplementary fig. 7, Supplementary Material online). This, along with Cygb being lost from all other sharks and rays, suggests that the Cygbs of the horn shark and white shark are pseudogenes.

**Figure 5:**
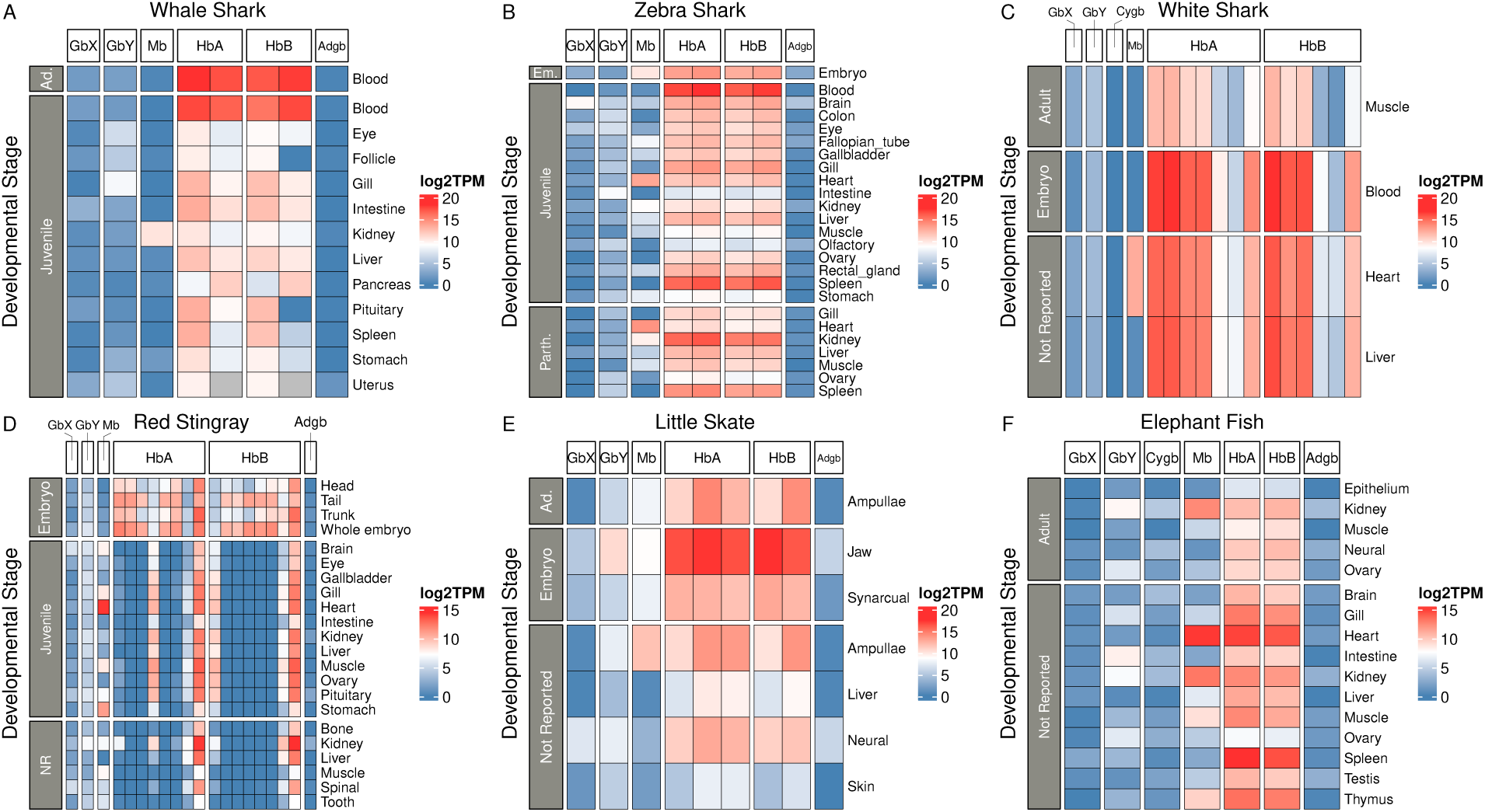
Expression of cartilaginous fish globin genes across tissues and developmental stages. Expression estimates were derived from log2-transformed TPM values assigned by Kallisto. Tissues and developmental stages were labeled according to information in the SRA run selector metadata. The full selection of taxa is shown in Supplementary fig. 7, Supplementary Material online. Em. = Embryo, Ad. = Adult, NR = Not Reported.

We see different patterns of expression in species with multiple α- and β-globin paralogs. In the red stingray, most α- and β-globin paralogs are highly expressed in early development (embryonic tissues), but the number of expressed paralogs diminishes in later developmental stages (fig. 5D). Expression data from the Giant manta and the Atlantic sting ray is consistent with this: a limited number of their α- and β-globin paralogs appear to be responsible for providing α- and β-globin chains for hemoglobin assembly (Supplementary fig. 7J-K, Supplementary Material online). However, sequence data from embryonic tissues in these species are lacking to confirm that more paralogs are expressed in earlier developmental stages. This does not seem to be the case in the white shark, where the proportional contribution of the different α- and β-globin paralogs does not seem to change much between embryos and adults (fig. 5C). Finally, in most species with less than 3 Hb paralogs per subfamily, all paralogs seem to be expressed, with small differences among them (Supplementary fig. 7, Supplementary Material online). The exception here is the lesser electric ray, where one of the three α-globin paralogs is expressed at much lower levels than the other two, and the whale shark, where one of two β-globin paralogs is expressed at much lower levels in follicle and pituitary tissues (Supplementary fig. 7N, Supplementary Material online).

### Ancestral Reconstruction and Evolution of the Globin Repertoire

The Adgb gene tree is concordant with the known organismal phylogeny, suggesting that this gene was present in the last common ancestor of cartilaginous fishes as single copy and has remained in this state in all descendant lineages. A strict reconciliation of the subtrees of the different globin paralogs in fig. 3 with the organismal tree in fig. 1 is straightforward for both α- globin and Cygb and implies that these genes were present in single copy state in both the last common ancestor of cartilaginous fishes and the last common ancestor of elasmobranchs, and that Cygb was secondarily lost multiple times independently (fig. 1). The cases of Ngb and GbE are also simple. Since they are absent from all the genomes screened, the most parsimonious inference is that they were also absent from their last common ancestor. This strict reconciliation would also imply the presence of multiple GbX1s and GbYs in the last common ancestor of cartilaginous fishes, and of multiple GbX1s, GbYs, Mbs, and β-globins in the last common ancestor of elasmobranchs. However, topology tests cannot distinguish between the best subtrees and those that forced Batoidea, Selachimorpha, and Holocephali sequences into monophyletic groups. Thus, we infer that these genes were also in a single copy state in the common ancestor of cartilaginous fishes and the common ancestor of elasmobranchs. In the case of GbY, as explained above, we also picked the constrained tree because we suspect that the result is a combination of unusually high rates of evolution in the bony fish portion of the tree and annotation inconsistencies.

Ignoring the elephant fish globins that were not present in the genome assemblies, the globin repertoire of the last common ancestor of cartilaginous fishes is inferred to have had single copies of Adgb, α- and β-globin, Cygb, Mb, GbY, GbX1 and GbX2. From there, the only change in the branch leading to the last common ancestor of Holocephali is the loss of GbX1. On the other side of the tree, the only change in the branch leading from the last common ancestor of cartilaginous fishes to the last common ancestor of elasmobranchs is the loss of GbX2, with the loss of Cygb in the last common ancestor of batoids, as well as in three different lineages in the Selachimorpha. The α- and β-globin subfamilies expanded independently in Selachimorpha and Batoidea. In the case of Selachimorpha, all these expansions lead to species-specific clades, whereas in the case of Batoidea, there are both species-specific expansions as well as older ones shared among different species. These expansions were most prominent in the α-globin gene family relative to the β-globin in most species, with the Giant manta providing the most extreme case with 20 copies of α-globin and 9 of β-globin.

### Evolution of the Globin Superfamily in Cartilaginous Fish and Bony Vertebrates

The globin repertoires of cartilaginous fish differ in key aspects relative to those of bony vertebrates. Cartilaginous fish lack Ngb, a globin which is present in almost all bony vertebrates and is heavily expressed in the brain and other neural tissues. Cygb is another highly conserved globin in bony vertebrates that, in cartilaginous fish, is either entirely absent, nonfunctional, or expressed at very low levels. By contrast, all surveyed cartilaginous fish retain a single copy of GbY, a gene that has been lost multiple times in bony vertebrates (Hoffmann et al., 2018; Schwarze et al., 2015; Storz et al., 2011).

Another notable difference between cartilaginous fish and bony vertebrates is the translocation of NPRL3 from its ancestral chromosomal location upstream of the α- and β-globin clusters and GbY (Supplementary fig. 6, Supplementary Material online). This opens interesting questions regarding the regulation of Hb expression in cartilaginous fish because regulatory elements in the introns of NPRL3 are linked to the regulation of where and when the different Hb genes of cyclostomes and gnathostomes are expressed, even though these hemoglobins have independent evolutionary origins. Either cartilaginous fishes have acquired novel regulatory elements, as with the translocated β-globin gene clusters of amniotes, or the same elements can act in trans conformation.

As for the Hb clusters themselves, it is interesting to note that tandemly arranged clusters of Hb genes seem to have expanded independently from a single gene multiple times in the evolution of jawed vertebrates (Hoffmann et al., 2018; Opazo et al., 2008, 2013; Opazo, Hoffmann, et al., 2015): in the expansions of Hb clusters of sharks (1), of batoids (2), and the β-globin cluster of amniotes (3). In most bony vertebrates, these expansions of the hemoglobin clusters fueled the emergence of hemoglobins with specialized biochemical roles, such as the Root-effect hemoglobins of fish, and of hemoglobins with specialized developmental roles, as the embryonic and fetal hemoglobins of mammals. The hemoglobin and myoglobin genes of lampreys are also arranged in clusters that are independent in origin relative to the hemoglobin clusters of jawed vertebrates (Schwarze et al., 2014), and also evolved specialized functional and developmental roles (Fago et al., 2018). Our analyses suggest that a similar pattern is present in both sharks and batoids, as there are differences in the time and tissues where the different hemoglobin genes are expressed (fig. 5, Supplementary fig. 7, Supplementary Material online).

In conclusion, our study shows that the early evolution of the globin gene superfamily of cartilaginous fishes involved the loss of several paralogs present in the last common ancestor of jawed vertebrates, and the differential retention of some paralogs relative to bony vertebrates.

Such patterns suggest that the division of labor among globin paralogs may have evolved differently in each of the two main lineages of jawed vertebrates.

## Materials and Methods

### Bioinformatic Searches

We screened the genomes of 20 cartilaginous fish species with genomes annotated using the NCBI RefSeq eukaryotic genome annotation pipeline (last accessed April 30^th^ 2025). This sample includes 11 shark species covering 5 of the 8 orders in the Selachimorpha, 8 batoid species including 4 species in order Myliobatiformes, a sawfish (order Rhinopristiformes), an electric ray (order Torpediniformes), and 2 skates (order Rajiformes); and a sole representative of the order Holocephali (Table 1, Supplementary Table 1, Supplementary Material online).

Reference genomes and their annotations (in GFF3 format) were downloaded from NCBI using BiomartR (Drost & Paszkowski, 2017). To increase representation of this last order, we manually annotated the globin repertoire of a ratfish and a rabbitfish, two additional species in the group for which there are reference genomes available that do not meet the RefSeq criteria. To search for globin sequences, we combined searches on protein databases seeded with the amino acid sequences using blastp, searches of translated nucleotide databases seeded with amino acid sequences using tblastn, and searches of nucleotide databases seeded with nucleotide sequences using blastn. Initial searches were seeded with a representative set of known globins from deuterostomes that included the full repertoire of globins of the elephant fish as reported by Opazo et al. 2015, and added the Adgb, α-globin, β-globin, Cygb, Mb, and Ngb of humans, plus the GbE, GbX, and GbY sequences from coelacanth. First, a protein blast database was made using publicly available gene models built by NCBI of all cartilaginous fish of interest. Next, globin seeds were aligned to the database using blastp with an evalue cutoff of 1e^-5^. Summary information was downloaded for the resulting hits using NCBI datasets v17.3.0 (O’Leary et al., 2024). NCBI GeneIDs were obtained for each protein, and only the longest isoform of each gene was used for a subsequent phylogenetic analysis to confirm their homology to the original globin seeds. To do this, the longest protein isoforms were aligned along with the globin search seeds using mafft‘s (v7.490) l-ins-i algorithm (Katoh & Standley, 2013), and a phylogenetic tree was inferred using IQ-Tree v2.0.7 under the JTT+F+R6 model (Minh et al., 2020; Nguyen et al., 2015). Support for the nodes was evaluated with 10,000 ultrafast bootstrap replicates (Hoang et al., 2018; Minh et al., 2013). This tree was only used to confirm homology among our initial searches, and a more comprehensive sampling was used for our final phylogenetic analysis (see subsequent methods). A representative set of the globins from cartilaginous fishes was added as seeds when searching for globins that were not found in our initial search.

### Data Curation

Because of differences in the protein domain structure, bioinformatic searches and phylogenetic analyses were conducted separately for Adgb, which includes a globin domain with the helices rearranged and fused to additional protein domains, and for all other globins, which include a single globin domain, with some terminal extensions. In the case of searches seeded with Adgb, we discarded sequences that only matched the calpain domain, which were retrieved in our initial searches.

We used similarity scores for the initial classification of the sequences into the different globin types. These initial classifications were adjusted with the results of phylogenetic and synteny analyses. Almost all genomes possess copies of Adgb, GbY, and Mb identified in the annotated genes through our blastp searches. Whenever these genes were absent from an assembly, we used alternative approaches to get the sequences. In the case of Adgb, because of the complexity of the gene, we only used tblastn searches, ordering the hits by position along the query, and patching the fragments, removing spurious overlaps. Even though the Adgbs of white shark, ratfish, and rabbitfish were found in the corresponding genomes, the hits cover >75% of the translated protein and are clearly incomplete. We infer that these genes are present and functional, but not fully covered in the current genome releases. We included the corresponding partial sequences in our phylogenetic analyses, but the resulting gene models need to be verified with additional data. In the case of Cygb, GbY, and Mb, we combined tblastn to identify the fragment where the gene was located, with blastn searches seeded with the Cygb, GbY, or Mb exons from a closely related cartilaginous fish to manually annotate the genes. When we searched the nucleotide databases using globin exons as seeds, we identified the first exon of GbY in the Lesser electric ray genome but found no trace of exons 2 or 3. Because GbY is present in all other cartilaginous fishes, we surveyed transcriptome data to look for this gene.

Doing so, we were able to obtain the full coding sequence of this GbY gene by assembling de novo a transcriptome from the lesser electric ray and matching transcripts to this exon (see Supplementary Information, Supplementary Material Online). We failed to find any matches that would correspond to exons 2 or 3 in the reference genome, so we assume this gene is present in this species but absent from the current assembly. Exons 2 and 3 from the red stingray GbY gene are annotated as part of the Luc7l gene in the current annotation, GeneID 140735596, so we reannotated the first exon by comparison with the GbY exons of the closely related Atlantic stingray. Using a similar approach, we found and exon of the Mb gene of the ratfish, but because there are no RNA-Seq data sets available for the ratfish we were unable to obtain the full coding sequence for the Mb gene. We assume that the Mb gene is present in the ratfish but is simply missing from the current assembly. Because of the short length of the fragment, we did not include the sequence in the phylogenetic analysis. Finally, in the case of GbX, we also used synteny information to classify the genes into either GbX1 or GbX2, following Gallagher and Macqueen (Gallagher & Macqueen, 2017), as we did in Hoffmann et al. (2021).

We found four genes annotated as globins of unusual lengths, such as α- and β-globins longer than 200 amino acids. These genes were blasted back to the cartilaginous fish protein database, excluding the source species. In the case of the 116985437 and 109935067 genes of thorny skate and whale shark, respectively, both are α-globins fused with a β-globin that were annotated as separate genes in previous genome releases; thus, the older annotations were used: ENSARAG00005009971 and ENSARAG00005009980 for the thorny skate and ENSRTYG00015001848 and ENSRTYG00015001843 for the whale shark. In addition, the 121288390 gene of the white shark and the 122539636 gene of the whitespotted bambooshark are also longer than our threshold. In these cases, the genes were discarded because they included non-globin-like domains.

### Phylogenetic analyses

For all phylogenetic analyses, we aligned the predicted amino acid sequences using the L-INS-i strategy from Mafft v7.505 (Katoh & Standley, 2013; Katoh & Toh, 2008). We then estimated phylogenetic relationships among the sequences using IQ-Tree version 3.0.1 (Wong et al., 2025), using the ModelFinder routine to estimate the best-fitting model of amino acid substitution (Kalyaanamoorthy et al., 2017), selecting the model chosen by the Bayesian Information Criterion, and using the resulting models in the searches (WAG+R6 for the full alignment). We evaluated support for the nodes with 10,000 replicates of the ultrafast bootstrap routine (Hoang et al., 2018). We compared competing phylogenetic hypotheses using constrained searches and the approximately unbiased topology test proposed by Shimodaira (2002) as implemented in IQ-Tree. A full description of the commands needed to replicate the analyses is available in Supplementary File 1, Supplementary Material online.

### Synteny analyses

We used a custom bash script to parse GFF annotations of the taxa used in this study and obtain genes flanking globins. Next, the longest protein isoforms corresponding to these genes were downloaded using NCBI datasets v17.3.0 (O’Leary et al., 2024). These proteins were aligned to the human reference proteome using blastp and were labeled with the gene name of the closest human blast hit. Syntenic regions were visualized using the gggenomes v1.0.1 package in R v4.4.2 (Hackl et al., 2024; R Core Team, 2020).

### Organismal Phylogeny and Divergence Dates

Phylogenetic relationships among cartilaginous fish species are based on the results of molecular studies (Aschliman et al., 2012; Naylor et al., 2012), and divergence times among the different groups correspond to the median time estimates provided by TimeTree v5 (Kumar et al., 2022).

### Structure Analyses

We investigated the structural differences between the α- and β-hemoglobin subunits of three cartilaginous fish, a great white shark (Selachimorpha), thorny skate (Batoidea), and elephant fish (Holocephali), and humans using the publicly available AlphaFold (Accessed May 1st, 2025). The protein sequences were uploaded to the web server, and a predictive 3D model was made from each of the protein sequences. The models were downloaded, and the CIF files were imported into ChimeraX v1.9 (Meng et al., 2023), where two of these models were overlayed on top of each other using the alignment feature.

### Gene Expression Analysis

RNA-Seq datasets from the cartilaginous fish used in our study were downloaded from the Sequence Read Archive (SRA) using the search term provided in the Supplementary Information, Supplementary Material online. Metadata for the resulting datasets was obtained from the SRA run selector (Supplementary Table 3, Supplementary Material Online). Sequence data was downloaded from the SRA using the SRAtoolkit commands “prefetch” and “fasterq-dump” (SRA Toolkit Development Team).

Reference transcriptomes for each species used in our study were downloaded from NCBI using Datasets (O’Leary et al., 2024). Separate Kallisto (v0.51.1) indices were made from each dataset using kallisto index, and reads were pseudo-aligned separately to the transcriptome indices using kallisto quant (Bray et al., 2016). Globin transcripts were extracted from Kallisto’s transcript abundance results and were labeled with their corresponding SRA datasets, GeneIDs, tissues, and developmental stages. Expression values for globin genes were derived from the sum transcripts per million (TPM) over all isoforms and log2 transformed for visualization. Manually annotated globin transcripts were added to their species’ transcriptome before indexing. Expression was visualized using the ComplexHeatmap v2.22.0 package in R (Gu et al., 2016; R Core Team, 2020).

## Acknowledgments

The authors thank the consortia and their personnel for releasing unpublished genome data and the associated metadata.

## Data Availability

All data used in this study were previously generated and can be found in NCBI’s RefSeq, GenBank, or sequence read archive (SRA) databases. Accession numbers for genomes used in this study are provided in Table 1. Metadata associated with globin sequences derived from bioinformatic searches is shown in Supplementary Table 1. Metadata related to SRA datasets used in the gene expression portion of this study are provided in Supplementary Table 3.

## References

Aschliman, N. C., Nishida, M., Miya, M., Inoue, J. G., Rosana, K. M., & Naylor, G. J. P. (2012). Body plan convergence in the evolution of skates and rays (Chondrichthyes: Batoidea). Molecular Phylogenetics and Evolution, 63(1). 10.1016/j.ympev.2011.12.012

Bray, N. L., Pimentel, H., Melsted, P., & Pachter, L. (2016). Near-optimal probabilistic RNA-seq quantification. Nature Biotechnology, 34(5), 525–527. 10.1038/nbt.3519

Brittain, T., & Wells, R. M. G. (1986). Characterization of the changes in the state of aggregation induced by ligand binding in the hemoglobin system of a primitive vertebrate, the hagfish Eptatretus cirrhatus. Comparative Biochemistry and Physiology -- Part A: Physiology, 85(4). 10.1016/0300-9629(86)90296-3

Burmester, T., Ebner, B., Weich, B., & Hankeln, T. (2002). Cytoglobin: A novel globin type ubiquitously expressed in vertebrate tissues. Molecular Biology and Evolution, 19(4). 10.1093/oxfordjournals.molbev.a004096

Burmester, T., Welch, B., Reinhardt, S., & Hankeln, T. (2000). A verteblrate globin expressed in the brain. Nature, 407(6803). 10.1038/35035093

Chong, K. T., Miyazaki, G., Morimoto, H., Oda, Y., & Park, S. Y. (1999). Structures of the deoxy and CO forms of haemoglobin from Dasyatis akajei, a cartilaginous fish. Acta Crystallographica Section D: Biological Crystallography, 55(7). 10.1107/S0907444999005934

Drost, H. G., & Paszkowski, J. (2017). Biomartr: Genomic data retrieval with R. Bioinformatics, 33(8). 10.1093/bioinformatics/btw821

Fago, A., Rohlfing, K., Petersen, E. E., Jendroszek, A., & Burmester, T. (2018). Functional diversification of sea lamprey globins in evolution and development. Biochimica et Biophysica Acta - Proteins and Proteomics, 1866(2). 10.1016/j.bbapap.2017.11.009

Fisher, W. K., Nash, A. W., & Thompson, E. O. P. (1977). Haemoglobins of the Shark, Heterodontus portusjacksoni III*. Amino Acid Sequence of the β -Chain. Australian Journal of Biological Sciences, 30(6). 10.1071/BI9770487

Fuchs, C., Burmester, T., & Hankeln, T. (2006). The amphibian globin gene repertoire as revealed by the Xenopus genome. Cytogenetic and Genome Research, 112(3–4). 10.1159/000089884

Gallagher, M. D., & Macqueen, D. J. (2017). Evolution and expression of tissue globins in ray-finned fishes. Genome Biology and Evolution, 9(1). 10.1093/gbe/evw266

Goodman, M. (1981). Globin evolution was apparently very rapid in early vertebrates: A reasonable case against the rate-constancy hypothesis. Journal of Molecular Evolution, 17(2). 10.1007/BF01732683

Goodman, M., Moore, G. W., & Matsuda, G. (1975). Darwinian evolution in the genealogy of haemoglobin. Nature, 253(5493). 10.1038/253603a0

Gu, Z., Eils, R., & Schlesner, M. (2016). Complex heatmaps reveal patterns and correlations in multidimensional genomic data. Bioinformatics, 32(18). 10.1093/bioinformatics/btw313

Hackl, T., Ankenbrand, M., van Adrichem, B., Wilkins, D., & Haslinger, K. (2024). gggenomes: effective and versatile visualizations for comparative genomics. https://arxiv.org/abs/2411.13556

Hoang, D. T., Chernomor, O., von Haeseler, A., Minh, B. Q., & Vinh, L. S. (2018). UFBoot2: Improving the Ultrafast Bootstrap Approximation. Molecular Biology and Evolution, 35(2), 518–522. 10.1093/molbev/msx281

Hoffmann, F. G., Opazo, J. C., Hoogewijs, D., Hankeln, T., Ebner, B., Vinogradov, S. N., Bailly, X., & Storz, J. F. (2012). Evolution of the globin gene family in deuterostomes: Lineage-specific patterns of diversification and attrition. Molecular Biology and Evolution, 29(7). 10.1093/molbev/mss018

Hoffmann, F. G., Opazo, J. C., & Storz, J. F. (2010). Gene cooption and convergent evolution of oxygen transport hemoglobins in jawed and jawless vertebrates. Proceedings of the National Academy of Sciences of the United States of America, 107(32). 10.1073/pnas.1006756107

Hoffmann, F. G., Opazo, J. C., & Storz, J. F. (2012). Whole-genome duplications spurred the functional diversification of the globin gene superfamily in vertebrates. Molecular Biology and Evolution, 29(1). 10.1093/molbev/msr207

Hoffmann, F. G., Storz, J. F., Kuraku, S., Vandewege, M. W., & Opazo, J. C. (2021). Whole-Genome Duplications and the Diversification of the Globin-X Genes of Vertebrates. Genome Biology and Evolution, 13(10). 10.1093/gbe/evab205

Hoffmann, F. G., Vandewege, M. W., Storz, J. F., & Opazo, J. C. (2018). Gene turnover and diversificationof the α-and β-GlobinGene families in sauropsid vertebrates. Genome Biology and Evolution, 10(1). 10.1093/gbe/evy001

Hoogewijs, D., Ebner, B., Germani, F., Hoffmann, F. G., Fabrizius, A., Moens, L., Burmester, T., Dewilde, S., Storz, J. F., Vinogradov, S. N., & Hankeln, T. (2012). Androglobin: A chimeric globin in metazoans that is preferentially expressed in mammalian testes. Molecular Biology and Evolution, 29(4). 10.1093/molbev/msr246

Ingram, V. M. (1961). Gene evolution and the hæmoglobins. Nature, 189(4766). 10.1038/189704a0

Kalyaanamoorthy, S., Minh, B. Q., Wong, T. K. F., von Haeseler, A., & Jermiin, L. S. (2017). ModelFinder: fast model selection for accurate phylogenetic estimates. Nature Methods, 14(6), 587–589. 10.1038/nmeth.4285

Katoh, K., & Standley, D. M. (2013). MAFFT multiple sequence alignment software version 7: improvements in performance and usability. Molecular Biology and Evolution, 30(4), 772– 780. 10.1093/molbev/mst010

Katoh, K., & Toh, H. (2008). Recent developments in the MAFFT multiple sequence alignment program. Briefings in Bioinformatics, 9(4). 10.1093/bib/bbn013

Kendrew, J. C., Bodo, G., Dintzis, H. M., Parrish, R. G., Wyckoff, H., & Phillips, D. C. (1958). A three-dimensional model of the myoglobin molecule obtained by x-ray analysis. Nature, 181(4610). 10.1038/181662a0

Keppner, A., Correia, M., Santambrogio, S., Koay, T. W., Maric, D., Osterhof, C., Winter, D. V., Clerc, A., Stumpe, M., Chalmel, F., Dewilde, S., Odermatt, A., Kressler, D., Hankeln, T., Wenger, R. H., & Hoogewijs, D. (2022). Androglobin, a chimeric mammalian globin, is required for male fertility. ELife, 11. 10.7554/eLife.72374

Keppner, A., Maric, D., Correia, M., Koay, T. W., Orlando, I. M. C., Vinogradov, S. N., & Hoogewijs, D. (2020). Lessons from the post-genomic era: Globin diversity beyond oxygen binding and transport. In Redox Biology (Vol. 37). 10.1016/j.redox.2020.101687

Kimura, M. (1981). Was globin evolution very rapid in its early stages?: A dubious case against the rate-constancy hypothesis. In Journal of Molecular Evolution (Vol. 17, Issue 2). 10.1007/BF01732682

Komiyama, N. H., Shih, D. T. B., Looker, D., Tame, J., & Nagai, K. (1991). Was the loss of the D helix in α globin a functionally neutral mutation? Nature, 352(6333). 10.1038/352349a0

Kugelstadt, D., Haberkamp, M., Hankeln, T., & Burmester, T. (2004). Neuroglobin, cytoglobin, and a novel, eye-specific globin from chicken. Biochemical and Biophysical Research Communications, 325(3). 10.1016/j.bbrc.2004.10.080

Kumar, S., Suleski, M., Craig, J. M., Kasprowicz, A. E., Sanderford, M., Li, M., Stecher, G., & Hedges, S. B. (2022). TimeTree 5: An Expanded Resource for Species Divergence Times. Molecular Biology and Evolution, 39(8). 10.1093/molbev/msac174

Kuraku, S. (2010). Palaeophylogenomics of the vertebrate ancestor - Impact of hidden paralogy on hagfish and lamprey gene phylogeny. Integrative and Comparative Biology, 50(1). 10.1093/icb/icq044

Meng, E. C., Goddard, T. D., Pettersen, E. F., Couch, G. S., Pearson, Z. J., Morris, J. H., & Ferrin, T. E. (2023). UCSF ChimeraX: Tools for structure building and analysis. Protein Science, 32(11). 10.1002/pro.4792

Minh, B. Q., Nguyen, M. A. T., & Von Haeseler, A. (2013). Ultrafast approximation for phylogenetic bootstrap. Molecular Biology and Evolution, 30(5), 1188–1195. 10.1093/molbev/mst024

Minh, B. Q., Schmidt, H. A., Chernomor, O., Schrempf, D., Woodhams, M. D., von Haeseler, A., & Lanfear, R. (2020). IQ-TREE 2: New Models and Efficient Methods for Phylogenetic Inference in the Genomic Era. Molecular Biology and Evolution, 37(5), 1530–1534. 10.1093/molbev/msaa015

Miyata, M., Gillemans, N., Hockman, D., Demmers, J. A. A., Cheng, J. F., Hou, J., Salminen, M., Fisher, C. A., Taylor, S., Gibbons, R. J., Ganis, J. J., Zon, L. I., Grosveld, F., Mulugeta, E., Sauka-Spengler, T., Higgs, D. R., & Philipsen, S. (2020). An evolutionarily ancient mechanism for regulation of hemoglobin expression in vertebrate red cells. Blood, 136(3). 10.1182/blood.2020004826

Müller, G., Fago, A., & Weber, R. E. (2003). Water regulates oxygen binding in hagfish (Myxine glutinosa) hemoglobin. Journal of Experimental Biology, 206(8). 10.1242/jeb.00278

Naoi, Y., Chong, K. T., Yoshimatsu, K., Miyazaki, G., Tame, J. R. H., Park, S. Y., Adachi, S. I., & Morimoto, H. (2001). The functional similarity and structural diversity of human and cartilaginous fish hemoglobins. Journal of Molecular Biology, 307(1). 10.1006/jmbi.2000.4446

Naylor, G. J. P., Caira, J. N., Jensen, K., Rosana, K. A. M., Straube, N., & Lakner, C. (2012). Elasmobranch Phylogeny: A Mitochondrial Estimate Based on 595 Species. In Biology of Sharks and Their Relatives: Second Edition. 10.1201/b11867-9

Nguyen, L.-T., Schmidt, H. A., von Haeseler, A., & Minh, B. Q. (2015). IQ-TREE: A Fast and Effective Stochastic Algorithm for Estimating Maximum-Likelihood Phylogenies. Molecular Biology and Evolution, 32(1), 268–274. 10.1093/molbev/msu300

O’Leary, N. A., Cox, E., Holmes, J. B., Anderson, W. R., Falk, R., Hem, V., Tsuchiya, M. T. N., Schuler, G. D., Zhang, X., Torcivia, J., Ketter, A., Breen, L., Cothran, J., Bajwa, H., Tinne, J., Meric, P. A., Hlavina, W., & Schneider, V. A. (2024). Exploring and retrieving sequence and metadata for species across the tree of life with NCBI Datasets. Scientific Data, 11(1), 732. 10.1038/s41597-024-03571-y

Opazo, J. C., Butts, G. T., Nery, M. F., Storz, J. F., & Hoffmann, F. G. (2013). Whole-genome duplication and the functional diversification of teleost fish hemoglobins. Molecular Biology and Evolution, 30(1). 10.1093/molbev/mss212

Opazo, J. C., Hoffmann, F. G., Natarajan, C., Witt, C. C., Berenbrink, M., & Storz, J. F. (2015). Gene turnover in the avian globin gene families and evolutionary changes in hemoglobin isoform expression. Molecular Biology and Evolution, 32(4). 10.1093/molbev/msu341

Opazo, J. C., Hoffmann, F. G., & Storz, J. F. (2008). Differential loss of embryonic globin genes during the radiation of placental mammals. Proceedings of the National Academy of Sciences of the United States of America, 105(35). 10.1073/pnas.0804392105

Opazo, J. C., Lee, A. P., Hoffmann, F. G., Toloza-Villalobos, J., Burmester, T., Venkatesh, B., & Storz, J. F. (2015). Ancient duplications and expression divergence in the globin gene superfamily of vertebrates: Insights from the elephant shark genome and transcriptome. Molecular Biology and Evolution, 32(7). 10.1093/molbev/msv054

Perutz, M. F., Rossmann, M. G., Cullis, A. F., Muirhead, H., Will, G., & North, A. C. T. (1960). Structure of Hæmoglobin: A three-dimensional fourier synthesis at 5.5-. resolution, obtained by X-ray analysis. In Nature (Vol. 185, Issue 4711). 10.1038/185416a0

Preston, A. E., Frost, J. N., Teh, M. R., Badat, M., Armitage, A. E., Norfo, R., Wideman, S. K., Hanifi, M., White, N., Roy, N. B. A., Babbs, C., Ghesquiere, B., Davies, J., Howden, A. J. M., Sinclair, L. V, Hughes, J. R., Kassouf, M., Beagrie, R., Higgs, D. R., & Drakesmith, H. (2025). Ancient genomic linkage of α-globin and Nprl3 couples metabolism with erythropoiesis. Nature Communications, 16(1), 2749. 10.1038/s41467-025-57683-z

Prothmann, A., Hoffmann, F. G., Opazo, J. C., Herbener, P., Storz, J. F., Burmester, T., & Hankeln, T. (2020). The Globin Gene Family in Arthropods: Evolution and Functional Diversity. Frontiers in Genetics, 11. 10.3389/fgene.2020.00858

R Core Team. (2020). R: A Language and Environment for Statistical Computing. https://www.r-project.org/

Roesner, A., Fuchs, C., Hankeln, T., & Burmester, T. (2005). A globin gene of ancient evolutionary origin in lower vertebrates: Evidence for two distinct globin families in animals. Molecular Biology and Evolution, 22(1). 10.1093/molbev/msh258

Schwarze, K., Campbell, K. L., Hankeln, T., Storz, J. F., Hoffmann, F. G., & Burmester, T. (2014). The globin gene repertoire of lampreys: Convergent evolution of hemoglobin and myoglobin in jawed and jawless vertebrates. Molecular Biology and Evolution, 31(10). 10.1093/molbev/msu216

Schwarze, K., Singh, A., & Burmester, T. (2015). The Full Globin Repertoire of Turtles Provides Insights into Vertebrate Globin Evolution and Functions. Genome Biology and Evolution, 7(7). 10.1093/gbe/evv114

Shimodaira, H. (2002). An approximately unbiased test of phylogenetic tree selection. Systematic Biology, 51(3), 492–508. 10.1080/10635150290069913

SRA Toolkit Development Team. SRA Toolkit. Https://Trace.Ncbi.Nlm.Nih.Gov/Traces/Sra/Sra.Cgi?View=software.

Storz, J. F. (2018). Hemoglobin: Insights into protein structure, function, and evolution. In *Hemoglobin: Insights into Protein Structure*, Function, and Evolution. 10.1093/oso/9780198810681.001.0001

Storz, J. F., Opazo, J. C., & Hoffmann, F. G. (2011). Phylogenetic diversification of the globin gene superfamily in chordates. In IUBMB Life (Vol. 63, Issue 5). 10.1002/iub.482

Storz, J. F., Opazo, J. C., & Hoffmann, F. G. (2013). Gene duplication, genome duplication, and the functional diversification of vertebrate globins. In Molecular Phylogenetics and Evolution (Vol. 66, Issue 2). 10.1016/j.ympev.2012.07.013

Tan, Y. Y., Kodzius, R., Tay, B. H., Tay, A., Brenner, S., & Venkatesh, B. (2012). Sequencing and Analysis of Full-Length cDNAs, 5′-ESTs and 3′-ESTs from a Cartilaginous Fish, the Elephant Shark (Callorhinchus milii). PLoS ONE, 7(10). 10.1371/journal.pone.0047174

Verde, C., De Rosa, M. C., Giordano, D., Mosca, D., De Pascale, D., Raiola, L., Cocca, E., Carratore, V., Giardina, B., & Di Prisco, G. (2005). Structure, function and molecular adaptations of haemoglobins of the polar cartilaginous fish Bathyraja eatonii and Raja hyperborea. Biochemical Journal, 389(2). 10.1042/BJ20050305

Vinogradov, S. N., Hoogewijs, D., Bailly, X., Arredondo-Peter, R., Guertin, M., Gough, J., Dewilde, S., Moens, L., & Vanfleteren, J. R. (2005). Three globin lineages belonging to two structural classes in genomes from the three kingdoms of life. Proceedings of the National Academy of Sciences of the United States of America, 102(32). 10.1073/pnas.0502103102

Wong, T. K., Ly-Trong, N., Ren, H., Baños, H., Roger, A. J., Susko, E., Bielow, C., De Maio, N., Goldman, N., Hahn, M. W., Huttley, G., Lanfear, R., & Quang Minh, B. (2025). IQ-TREE 3: Phylogenomic Inference Software using Complex Evolutionary Models. EcoEvoRxiv. 10.32942/X2P62N

