## Supplementary Information for "A Deep Dive into the Globin Superfamily of Sharks, Skates, and Rays: Contrasting patterns of gene loss and retention relative to bony vertebrates"

Search term for NCBI SRA (Conducted on May 7, 2025)

(("Chiloscyllium plagiosum"[Organism] OR Chiloscyllium plagiosum[All Fields]) OR ("Pristis pectinata"[Organism] OR Pristis pectinata[All Fields])) OR Amblyraja radiata OR Carcharodon carcharias OR Callorhinchus milii OR Hemiscyllium ocellatum OR Rhincodon typus OR Leucoraja erinacea OR Hypanus sabinus OR Stegostoma tigrinum OR Heptranchias perlo OR Narcine bancroftii OR Pristiophorus japonicus OR Scyliorhinus canicula OR Mobula hypostoma OR "Scyliorhinus torazame"[Organism] OR "Scyliorhinus torazame"[All Fields]) OR ("Mobula birostris"[Organism] OR "Mobula birostris"[All Fields]) OR ("Hemitrygon akajei"[Organism] OR "Hemitrygon akajei"[All Fields]) OR ("Chiloscyllium punctatum"[Organism] OR "Chiloscyllium punctatum"[All Fields]AND ("biomol rna"[Properties] AND "library layout paired"[Properties])

Datasets removed after search

SRR12550959 ignored because it was ncRNAseq

SRR21023205-SRR21023210 ignored because it was amplicon

SRR27125045-SRR27125050 ignored because it was Assay type "other" with no information.

SRR31783813-SRR31783816 ignored because it was Chipseq

Lesser Electric Ray Transcriptome Assembly

SRA datasets SRR29142957 and SRR29142959 derived from lesser electric ray (LER) tissues were concatenated together and read depth was normalized to a depth of 10,000 using Trinity’s insilico_read_normalization.pl script (Grabherr et al., 2011). These reads were then provided to Trinity via their docker container with the –no_normalize_reads option on. The LER transcriptome was made into a nucleotide blast database, and a partial GbY sequence derived from the LER reference genome (exon 1 of the putative GbY) was alinged to the transcriptome using blastn. From the resulting blast hits, we found at transcript of length 1,520 nt, encoding for a 160 aa protein with high homology to other catilaginous fish GbYs.
