## Supplementary Table 2 for "A Deep Dive into the Globin Superfamily of Sharks, Skates, and Rays: Contrasting patterns of gene loss and retention relative to bony vertebrates"

Supplementary Table 3. Results of topology tests where we compared the maximum likelihood tree (ML) for each paralog with one where the Holocephali, Batoidea and Selachimorpha sequences were constrained to be monophyletic. All tests were run in IQ-Tree 3.0.1 for Linux 64-bit (Wong et al. 2025)

| Globin  paralog | logL of  ML tree | logL of  constrained tree | ΔL | p-AU |
| --- | --- | --- | --- | --- |
| β-globin | -7094.6 | -7095.4 | 0.8 | 0.28 |
| Mb | -3734.7 | -3748.5 | 12.1 | 0.14 |
| GbY | -3181.8 | -3191.1 | 9.3 | 0.03 * |
| GbX | -6250.3 | -6255.1 | 2.0 | 0.34 |

ΔL: difference in likelihood score

p-AU: p-value of approximately unbiased (AU) test (Shimodaira, 2002)

References

Shimodaira H. 2002. An approximately unbiased test of phylogenetic tree selection. *Syst Biol*. 513:492–508.

T.K.F. Wong, N. Ly-Trong, H. Ren, H. Banos, A.J. Roger, E. Susko, C. Bielow, N. De Maio, N. Goldman, M.W. Hahn, G. Huttley, R. Lanfear, B.Q. Minh. 2025. IQ-TREE 3: Phylogenomic Inference Software using Complex Evolutionary Models. Submitted, https://doi.org/10.32942/X2P62N.
