## Supplementary Figure 1 for "A Deep Dive into the Globin Superfamily of Sharks, Skates, and Rays: Contrasting patterns of gene loss and retention relative to bony vertebrates"

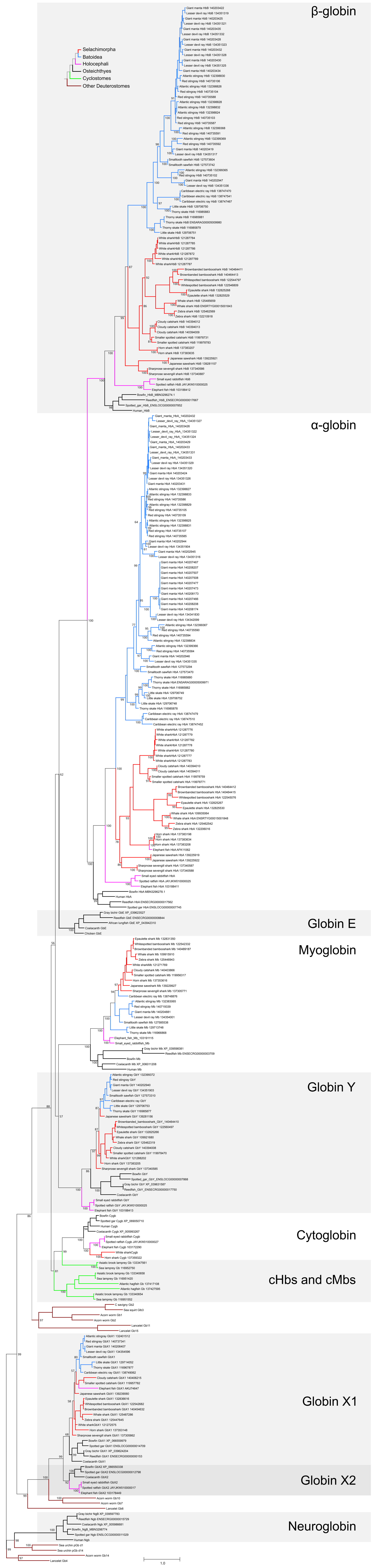


**Supplementary Figure 1**. Maximum likelihood phylogram depicting relationships among the single-domaing globin genes of cartilaginous fishes, with bony vertebrate and deuterostome sequences included for context. Numbers corresp-ond to ultrafast bootstrap values. Branches are colored according to the tree in the inset. Genes marked with asterisks are included in the phylogeny for comparative purposes but they are not found in the current genome
