## Supplementary Figure 2 for "A Deep Dive into the Globin Superfamily of Sharks, Skates, and Rays: Contrasting patterns of gene loss and retention relative to bony vertebrates"

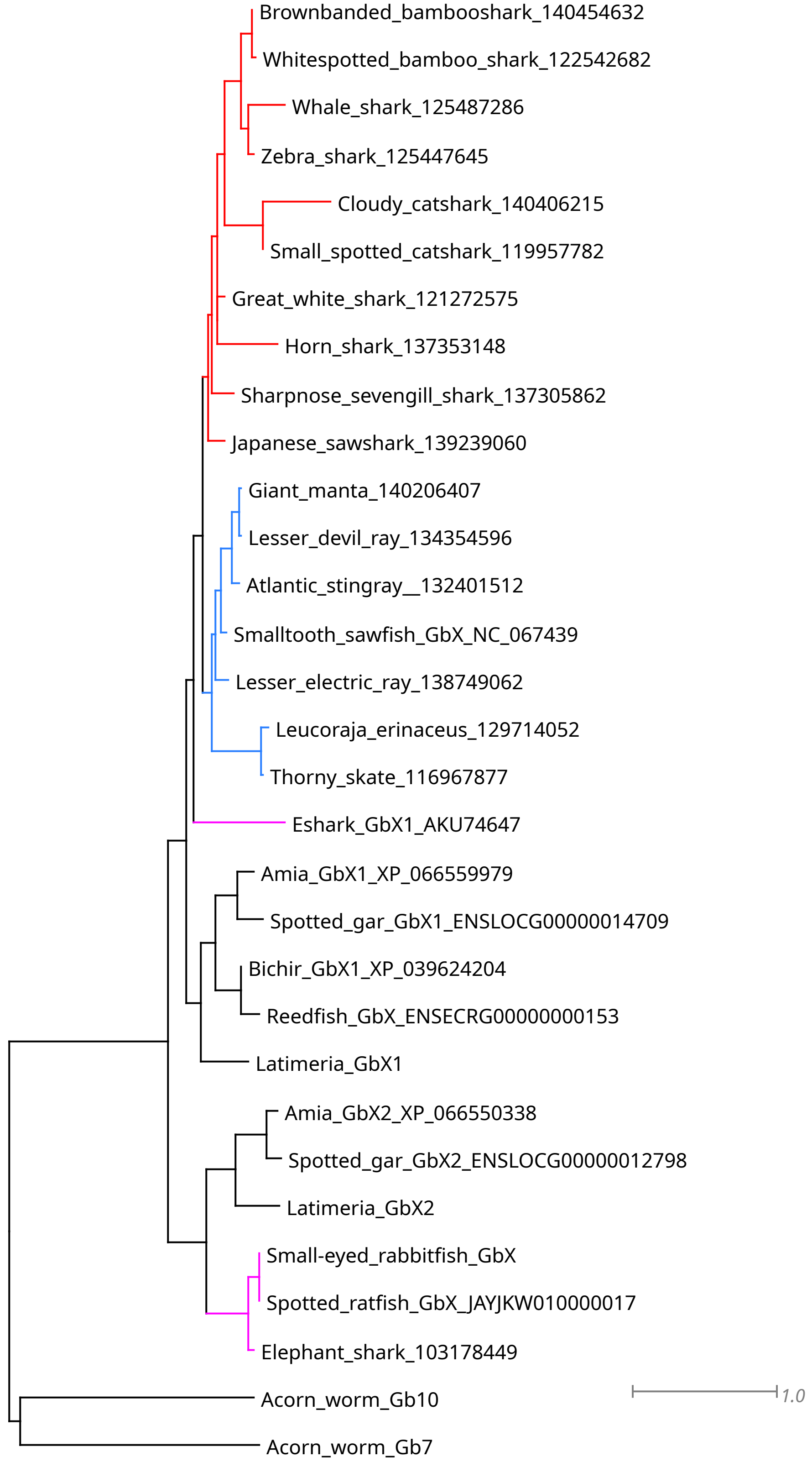


Supplementary Figure 2. Maximum likelihood phylogram depicting relationships among the GbX genes of cartilaginous fishes forcing GbX1 and GbX2 to be reciprocally monophyletic while constraining the cartilaginous fish GbX1s of Batoidea and Selachimorpha to be monophyletic. Shark branches in red, batoid branches in blueand holostei branches in fucsia.
