## Supplementary Figure 3 for "A Deep Dive into the Globin Superfamily of Sharks, Skates, and Rays: Contrasting patterns of gene loss and retention relative to bony vertebrates"

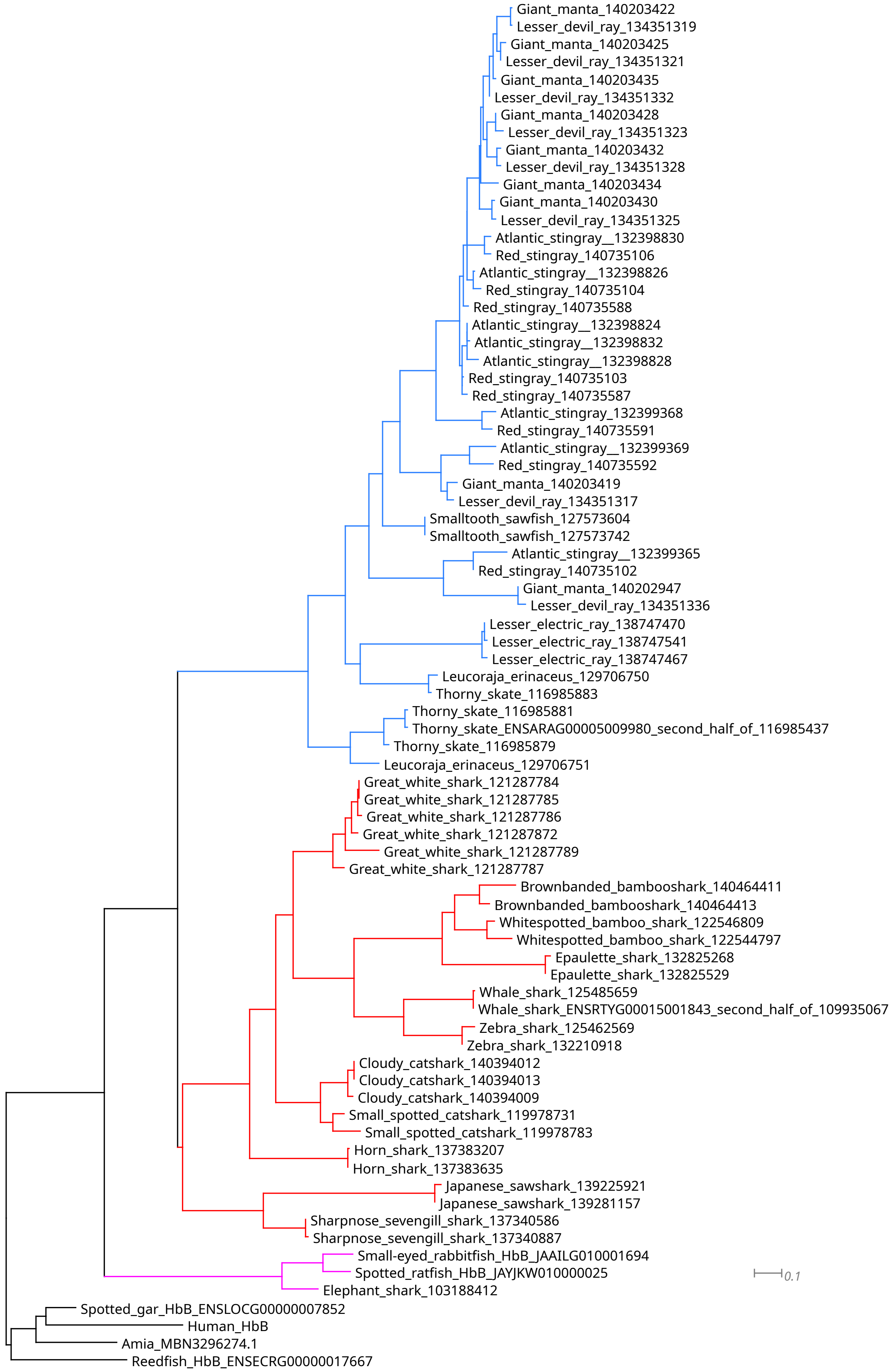


Supplementary Figure 3. Maximum likelihood phylogram depicting relationships among the β-globin genes of cartilaginous fishes forcing forcing each of Holocephali, Batoidea and Selachimorpha sequences to be monophyletic. Shark branches in red, batoid branches in blue and holostei branches in fucsia.
