## Supplementary Figure 4 for "A Deep Dive into the Globin Superfamily of Sharks, Skates, and Rays: Contrasting patterns of gene loss and retention relative to bony vertebrates"

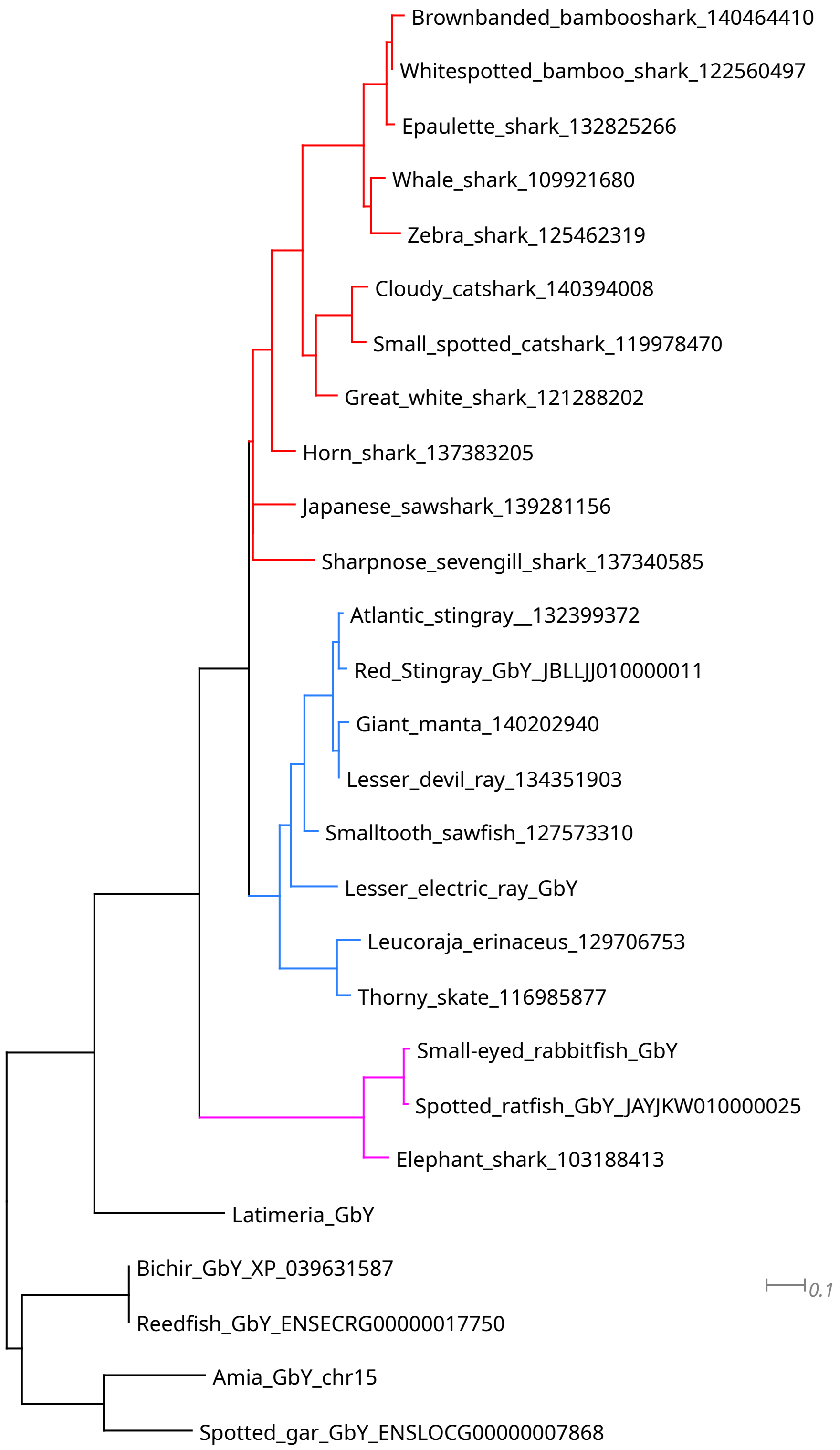


Supplementary Figure 4. Maximum likelihood phylogram depicting relationships among the GbY genes of cartilaginous fishes forcing forcing each of Holocephali, Batoidea and Selachimorpha sequences to be monophyletic. Shark branches in red, batoid branches in blue and holostei branches in fucsia.
