## Supplementary Figure 7 for "A Deep Dive into the Globin Superfamily of Sharks, Skates, and Rays: Contrasting patterns of gene loss and retention relative to bony vertebrates"

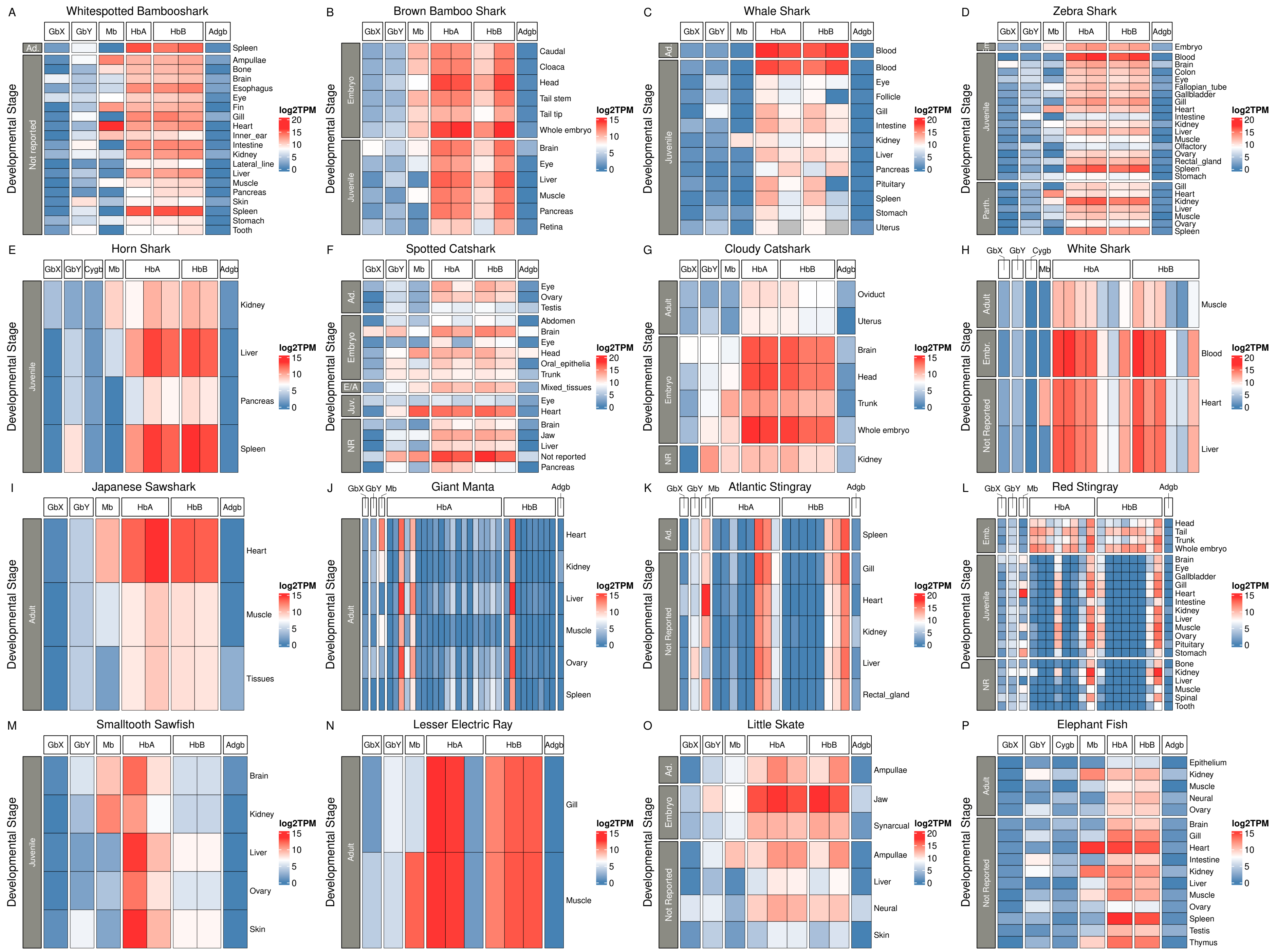


Supplementary Figure 7. Globin expression of cartilaginous fish across tissues and developmental stages. Expression estimates were derived from log2 transformed TPM values assigned by Kallisto. Tissues and developmental stages were labeled according to information in the SRA run selector metadata. Em. = Embryo, Ad. = Adult, NR = Not Reported, E/A = Embryo/Adult.
