## Supplementary Figure 6 for "A Deep Dive into the Globin Superfamily of Sharks, Skates, and Rays: Contrasting patterns of gene loss and retention relative to bony vertebrates"

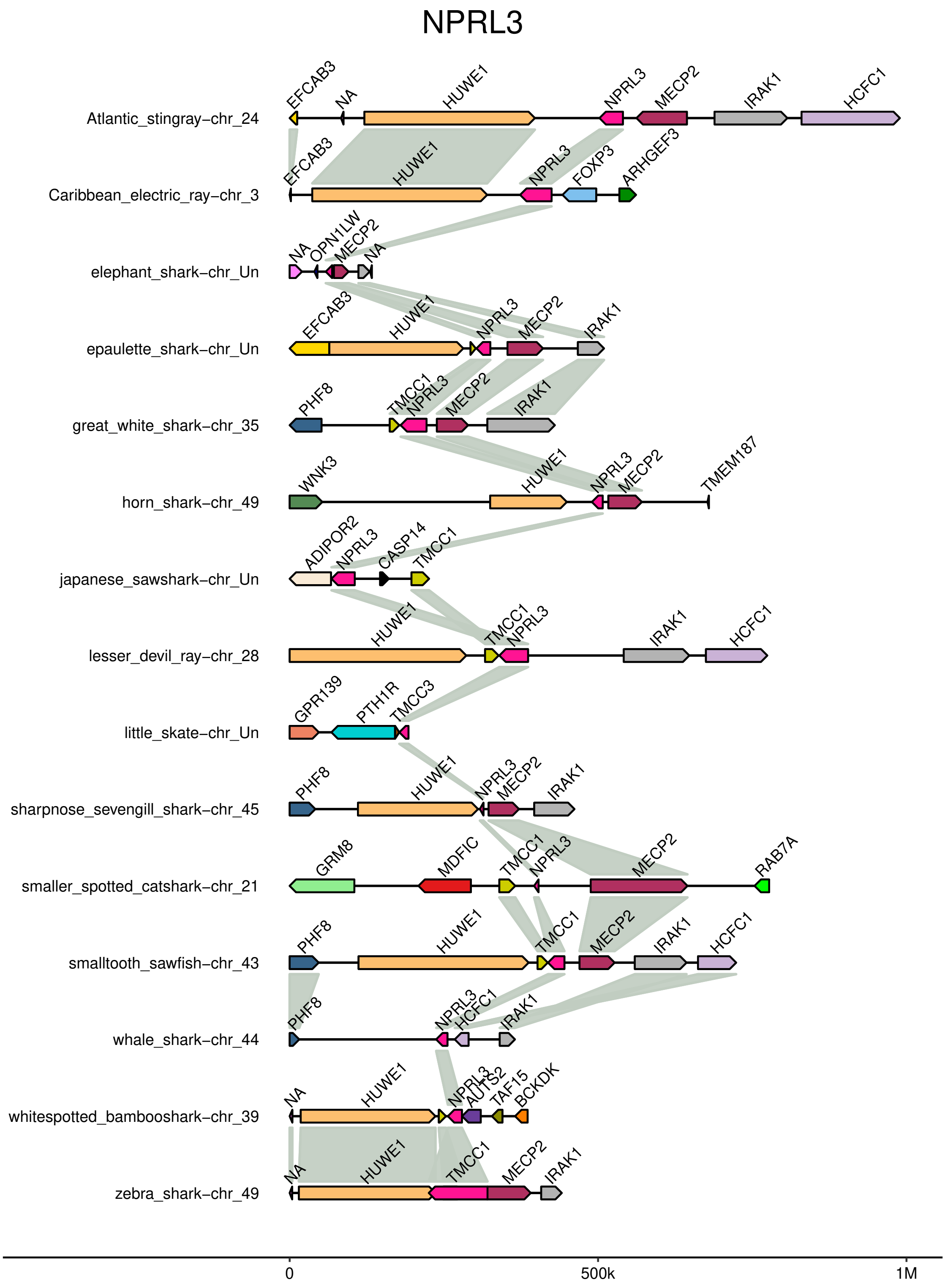


Figure S7. Synteny plot of cartilaginous fish NPRL3. Gray blocks connect putatitve orthologs between species
